# Machado: open source genomics data integration framework

**DOI:** 10.1101/2020.05.08.084731

**Authors:** Mauricio de Alvarenga Mudadu, Adhemar Zerlotini

## Abstract

**Background:** Genome projects and multiomics experiments generate huge volumes of data that must be stored, mined and transformed into useful knowledge. All this information is supposed to be accessible and, if possible, browsable afterwards. Computational biologists have been dealing with this scenario for over a decade and have been implementing software libraries, toolkits, platforms, and databases to succeed in this matter. The GMOD’s (Generic Model Organism Database project) biological relational database schema, known as *Chado*, is one of the few successful open source initiatives, it is widely adopted and many softwares are able to connect to it.

**Results:** We have been developing an open source software named *Machado* (https://github.com/lmb-embrapa/machado), a genomics data integration framework implemented in *Python*, to enable research groups to both store and browse, query, and visualize genomics data. The framework relies on the *Chado* database schema and, therefore, should be very intuitive for current developers to adopt it or have it running on the top of already existing databases. It has several data loading tools for genomics and transcriptomics data and also for annotation results from tools such as *BLAST*, *InterproScan*, *OrthoMCL* and *LSTrAP*. There is an API to connect to *JBrowse* and a web browsing visualisation tool is implemented using *Django Views and Templates*. The *Haystack* library integrated with the *ElasticSearch* engine was used to implement a google-like search i.e. single auto-complete search box that provides fast results and incremental filters.

**Conclusion:** *Machado* aims to be a modern object-relational framework that uses the latests *Python* libraries to produce an effective open source resource for genomics research.

## BACKGROUND

The technological advances for biological research regarding genomic sequencing, phenotype prediction and re-engineering of living systems have led to the creation of large volumes of data that must be stored, mined and transformed into useful knowledge. These technological advances of the omic approaches, including genomics, transcriptomics, proteomics and metabolomics, have great impact in several areas of knowledge including agriculture, specially for integrating big data analysis into animal and plant breeding programs.

Omics enables a systems biology approach toward understanding the complex interactions between genes, proteins, and metabolites within the resulting phenotype (Van Emon, 2016). Omics data integration offers the potential to increase the productivity and sustainability in crop and livestock production.The challenges are diverse but are usually composed of identifying genetic variation that derive desirable traits that can drive genomic prediction, performing precise genome editing/engineering (e.g.: using CRISPR-CAS systems for the induction of mutations or disruptions in the genome), identifying molecular targets for developing vaccines to diseases/plagues, and probably others (Huang et al., 2017).

All these novel genomic information, specially those from genome projects and multiomics experiments (transcriptomics, proteomics, etc.) is supposed to be accessible and, if possible, browseable afterwards. Even further, another great challenge is to integrate data from different organisms and projects for analysis and mining of data. Plant and animal trait data are typically generated by diverse, costly and time-consuming experiments, and thus can hugely benefit from increased data sharing and integration (Leonelli et al., 2017). Bioinformaticians and computational biologists have been dealing with this scenario for over a decade now and have implemented (and are still implementing) a collection of software libraries, toolkits, platforms, databases and data warehouses in this regard.

Although public wide databases exist, research groups still struggle to store and analyze data with local resources and expertise. The GMOD, Generic Model Organism Database project, is currently the initiative that most advanced in producing a “collection of open source software tools for managing, visualizing, storing and disseminating genomic data” (GMOD, 2020). Its biological relational database schema named Chado (Mungall & Emmert, 2007) is widely adopted and many softwares are able connect to it eg. Jbrowse (Skinner et al., 2009), Gbrowse (Stein et al., 2002), Apollo (Lee et al., 2013), Intermine (Kalderimis et al., 2014), and Tripal (Spoor et al., 2019).

The development of a front-end for Chado named Tripal, based on the Drupal CMS, facilitated the construction and publication of genomic databases (Tripal, 2020), although historically PHP is barely used in bioinformatics. For instance, BioPHP, a collection of open source PHP code with a number of bioinformatics tools latest release is from 2003 (“BioPHP,” 2020).

“Python has arguably become the de facto standard for exploratory, interactive, and computation-driven scientific research” (Millman & Aivazis, 2011). Ranking first in the 2019 IEEE Spectrum top programming languages (Cass, 2019), Python has a vast collection of libraries and modules for bioinformatics eg. BioPython (Cock et al., 2009), PyVCF, and PySAM; and data science eg. SciPy, NumPy, pandas, and Matplotlib. Coupled with Django, one of the most mature and feature-rich web application framework for Python (Django, 2020), developers are able to produce fast, secure, and scalable softwares.

The Embrapa’s Bioinformatics Multi-user Laboratory began to develop an open source software called Machado that has a Django Model to connect to Chado, thus avoiding extra efforts to make data compatible to the database schema.

The Chado database schema enables us to integrate different data types using controlled vocabularies and ontologies. For example, the Sequence Ontology (Eilbeck et al., 2005), a collection of sequence feature types, is used for typing features in the sequence module of Chado. Therefore, every biological component and its relationships are formally described, allowing the identification of exons that are ‘part_of’ a gene or transcripts that are ‘part_of’ a gene. Additionally, the Gene Ontology (Gene Ontology Consortium, 2004) enables us to functionally characterize the biological components regarding to their molecular function, cellular localization and the biological process they are involved with.

The Django framework provides practical means to build visualization tools and APIs (Application Programming Interface) to assist software developers to deal with multiple genomic data sources for building seamless, interoperable applications. The API framework is a set of clearly defined methods of communication between various software components. The data standardization across different research groups, coupled with the API framework, will facilitate future collaborations with data scientists in order to explore the data sets even further.

We intend to provide a Python framework to store diverse biological data, make complex queries and visualize results for this project. But even further, we hope to broaden the Machado software usability and provide the community with a powerful, simple, open source software that could be used by other scientific groups in projects of many ranges of complexities.

## METHODS

### The database model

The *Machado* software (https://github.com/lmb-embrapa/machado) was developed and tested using Python3 and Django 2.2.10. The first step was to create an object-relational model for the GMOD Chado database schema 1.31, using the Django inspectdb tool and a custom Python script that is available within the Machado repository. The resulting model integrates the Machado package and is used solely to connect to a Chado database, not to create the tables. During a new Machado installation, the original GMOD Chado schema SQL file is used to create the database instance. Therefore, users should be able to provide an already populated database instance if desired.

### The data loading tools

The data loading tools were developed following the interface segregation principle, in which, the classes related for ETL (extract, transform, and load) are independent from the classes related to the user interface. This design pattern makes it possible to implement diverse user interfaces (command-line, web, or API) while preserving the ETL classes.

The user interface was implemented using the Django management tool, a command-line management system to execute database related tasks in a standard manner. The management tool, together with a few Python libraries, allowed us to implement an effective set of data loading commands capable of loading data files using multi-threading and providing users with real-time progress monitoring.

The ETL classes benefit from well-established bioinformatics libraries to handle the genomics data files, namely, BioPython to load FASTA, GFF, Blast, and InterproScan; obonet to load ontology files; and bibtexparser to load Bibtex files.

### The web interface

The Machado web interface was developed using the Django views and templates, which is a convenient way to generate HTML dynamically. The core set of web pages is constituted of a search form, a search results page, and a feature page.

The search form is embedded in the page header in order to allow users to conveniently search for features at any time. The search form contains autocomplete capabilities to help users to validate keywords i.e. to identify keywords that are stored in the database and its correct spelling. Once a query is executed, the search form redirects the web browser to the results page, which contains basic information of every feature that meets the search criteria. The search results are paginated and this page also provides search results filtering, text download, sequence download, column sorting, and selection of the number of records per page.

Every feature has an hyperlink that leads to the feature page, which contains every information stored in the database related to the feature itself eg. genes, transcripts, and proteins. Currently, it’s set to show ID, description, relationships, genomic location, functional annotation, similarity analysis results, co-expression networks, groups of orthologs, expression data, sequence, and related publications. The genomic location is rendered to an embedded JBrowse instance in order to enhance the users experience. It shows the genomic region in detail, the gene structure with its exons, and UTRs, and allows to zoom in/out to identify adjacent genes.

### The search engine

The Machado search and filter components are powered by the ElasticSearch engine that is invoked via the Haystack library. This approach allowed us to create a single search box that autocompletes the user keywords and finds matches throughout the whole database. The search results can be filtered by organism, type, orthology, coexpression, annotation, expression biomaterial, and expression treatment. This filtering component was implemented using the ElasticSearch faceted navigation which allow users to filter through vast data sets by checking or unchecking filters. Both the search and the indexing components are invoked via the Haystack library, which significantly simplified the software programming process since it uses familiar Django syntax, rather than native ElasticSearch coding. The Haystack library is able to directly connect to the Django model and retrieve data to a few search engines, such as, ElasticSearch, and Solr. The Machado repository contains the specific code to generate the search index based on the Chado schema. Therefore, after having all the data loaded up, the user will simply build the search index using a specific command in the Django management tool.

### Availability of Supporting Source Code and Requirements

Project name: Machado

Project home page: https://github.com/lmb-embrapa/machado

Operating system(s): Platform independent

Programming language: Python 3

Other requirements: Latest Pyhton version, Python development libraries

License: e.g. GNU GPL

bio.tools id: machado

RRID: SCR_018428

The complete source code is available in GitHub (https://github.com/lmb-embrapa/machado) and the tests are executed routinely, triggered by every new code commit by Travis-CI. Extensive testing and code reviews ensure the software is fully functional upon new installations. There’s also detailed instructions hosted at Read The Docs (https://machado.readthedocs.io) on how to install, load data, and set up the user interface.

## OVERVIEW

*Machado* is a Django instance that provides data management, visualization, and searching functionalities to Chado databases. The resulting object-relational framework enables users, not only to set up a local instance containing data regarding their organisms of interest, but also to develop all sorts of tools by accessing the open source code.

The data loading tools are currently available via the Django management interface. Such tools were develop to load data from the most common bioinformatics file formats to the Chado database. This implementation provides users with commands capable of loading data files using multi-thread and real-time progress monitoring. Developers are welcome to create or propose new data formats to future versions.

Machado also provides users with an out-of-the-box solution to browse the data, which is able to display every data loaded using the current data loading tools. This web interface is fully customizable in order to encourage the development of new solutions or the connection to analysis tools.

The current version can be tested at a demonstration website which contains genomics data from five plants and is available at https://www.machado.cnptia.embrapa.br. This interface aims to simplify data searching by providing users with a single search box to query all the data. The user would open the web page and type a given term (Figure 1-A) to have instant access to similar valid keywords provided by the autocomplete feature. After typing the keywords and submitting the form, the user is redirected to the search results page (Figure 1-B). Such page contains summarized information about genes, transcripts, and proteins from all the organisms and filtering boxes at the left section which enables selecting features by specific criteria, such as organism, type, orthology, coexpression, and annotation. It also provides column sorting, control of the number of results, and download of the table or the sequences (Figure 1-C).

**Figure 1:**
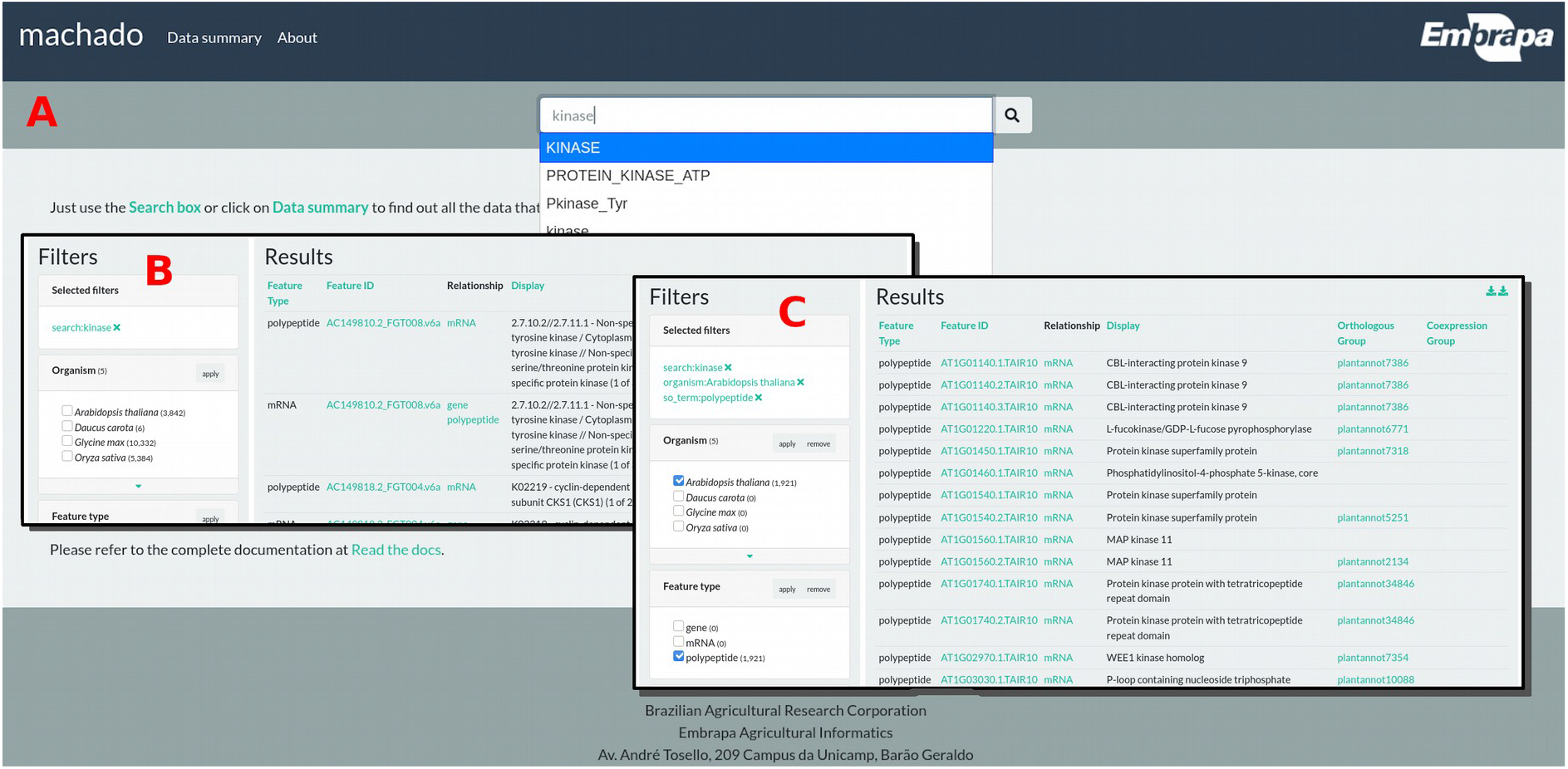
A) the Machado demonstration site home page, the search form, and autocomplete component; B) the search results for the term ‘kinase’; C) the search results for the term ‘kinase’, filtered by organism ‘Arabidopsis thaliana’, and and feature type ‘polypeptide’.

The search results columns contain hyperlinks to the feature itself (eg. protein), its related feature (eg. mRNA), orthologous groups, or coexpression groups. By clicking in a Hyperlink of a feature the user is redirected to the feature page that contains detailed information about it (Figure 2).

**Figure 2:**
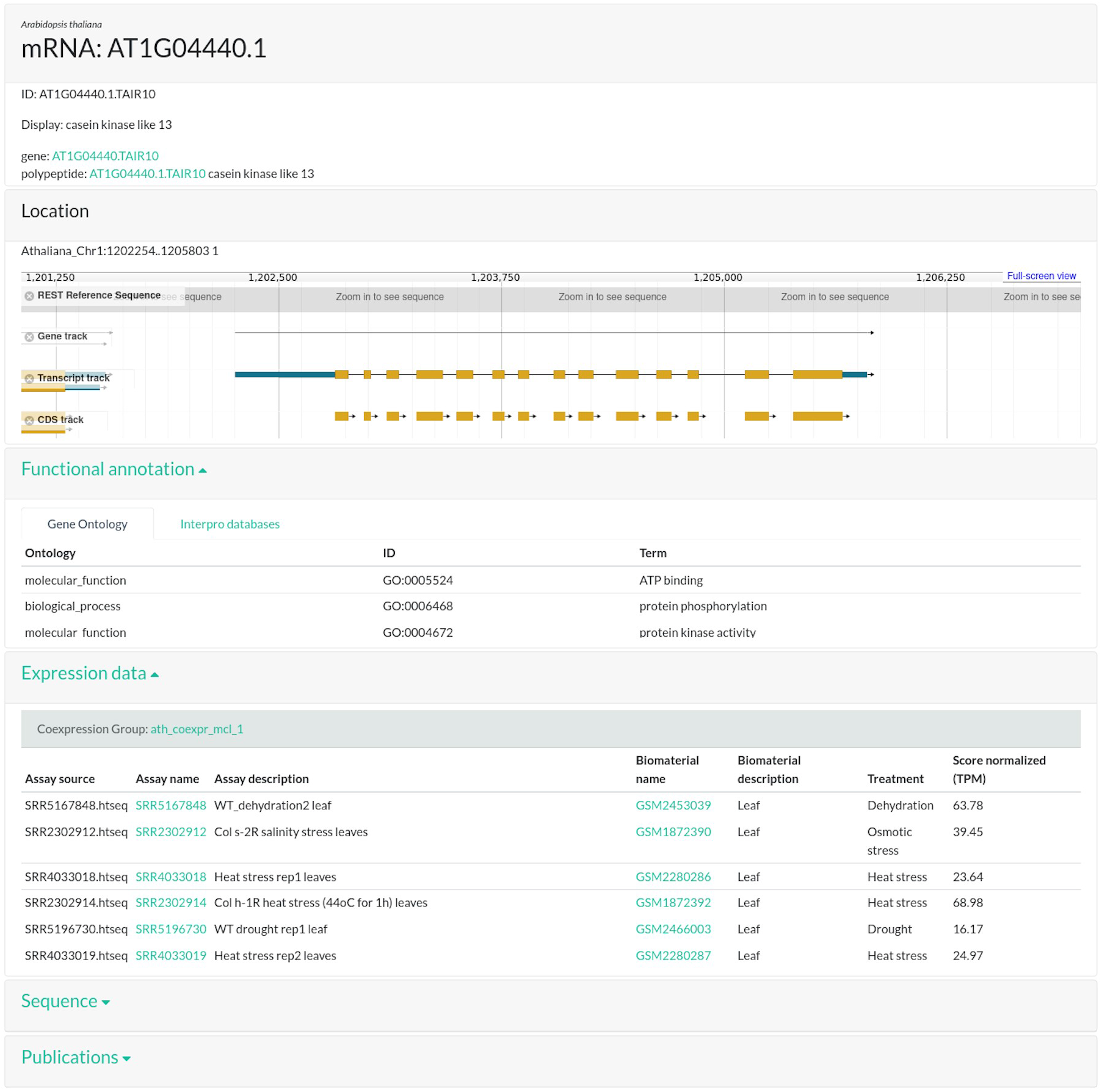
Feature page

The feature page is organized in cards that can be collapsed to facilitate the visualization of specific information. The first card contains information about the feature itself, such as the organism, IDs, related features, and text annotation. If the feature is mapped to the genome, the next card will show the genomic location and an embedded genome browser powered by JBrowser. The following cards contain detailed information about functional annotation, orthologs groups, coexpression groups, sequence, and publications.

## DISCUSSION AND CONCLUSION

Although Machado aims to simplify the genomics data integration, it requires considerable understanding of the data that’s being loaded as well as of the computational tools. The GMOD Chado database schema relies upon ontologies to establish the relationship among data types and, therefore, the data must be loaded in a particular order to ensure the parent data is always loaded in advance. The user must observe the file format specifications and ascertain the feature IDs are consistent throughout the files. In regard to the computational tools, it’s required to carry out systems administration tasks related to software installation and configuration, users permissions, and hardware requirements evaluation. There are other frameworks to integrate genomic data such as Intermine or Tripal that are in more advanced stages of development, but nevertheless users will have to go through the same laborious tasks described above.

However, the development of Machado was proven to be very fortunate once we started taking advantage of the Python libraries. For instance, Biopython enabled us to parse several file formats effectivelly and Haystack provided us very fast search and filtering capabilities. The single search box with autocomplete and faceting capabilities is arguably unprecedented among the open source frameworks available. The Python library repository is vast and, therefore, there is much to expand on future releases. For example, Machado can be used not to only host genomics data but also to enable the development of specific tools, such as, PlantAnnot (http://www.machado.cnptia.embrapa.br/plantannot), to identify and annotate genes of interest. There’s extensive documentation within the Machado repository and users are welcome to contact us and propose documentation updates.

The machado software is a modern open source python framework that intends to store, integrate, query, and visualize multiomics data and also to be fast and easy to use. The software is public and everyone can download, use and collaborate by proposing improvements and submitting code.

## ACKNOWLEDGEMENTS

Many thanks for Embrapa multiuser bioinformatics laboratory (LMB - Laboratório Multiusuário de Bioinformática da Embrapa) and Embrapa Agricultural Informatics (Embrapa Informática Agropecuária) for all the support.

